# Development of Ingenui-T, a Novel Vein-to-Vein Solution for Rapid Autologous CAR T-Cell Manufacturing Starting From Whole Blood, for the Treatment of Autoimmune Diseases

**DOI:** 10.1101/2024.01.24.576713

**Authors:** Daniel Anaya, Brandon Kwong, Soo Park, Sunetra Biswas, Jeevitha Jeevan, Madison Strobach, Nicole Khoshnoodi, Ames Register, Timothy D Klasson, Santiago Foos-Russ, Jennifer Zeng, Jesus Banuelos, Candice Gibson, Jazmin Bravo, Jeanne Flandez, Tom Van Blarcom, Karen Walker

**Author notes:** Address correspondence to: Name: Sunetra Biswas, Address: 5980 Horton St Suite 550, Emeryville, CA 94608. These authors contributed equally to the content (Daniel Anaya, Brandon Kwong, Soo Park).

## Abstract

Apheresis, a conventional starting point for manufacturing chimeric antigen receptor (CAR) T-cell therapy, poses challenges due to the length and invasiveness of the procedure, the high demand for and limited quantity of apheresis beds, and additional resource constraints at collection centers. Furthermore, traditional CAR T-cell manufacturing often involves extended cell culture periods, leading to a final product that has progressed through the differentiation process and contains a higher frequency of cells with phenotypes that are indicative of lower functionality or potency.^1^ Here, we show that anti-CD19 CAR T-cells manufactured from fresh whole blood with minimal ex vivo expansion using Ingenui-T exhibit comparable or superior CAR-mediated and CD19-dependent functional activity compared to CAR T-cells manufactured from cryopreserved leukapheresis material following the a conventional manufacturing process. Anti-CD19 CAR T-cell production, manufactured from whole blood in less than 3 days of in vitro culture using Ingenui-T, yielded an average of about 40 million cells per 100 mL with high CD3 purity and viability. Furthermore, the final product was composed of cells with less differentiated phenotypes and sustained cytotoxic activity against CD19^+^ target cells at a lower dose than conventionally manufactured CAR T cells in preclinical in vitro assays. Ingenui-T is an innovative vein-to-vein solution, aimed at enhancing the patient experience, feasibility, and accessibility of CAR T-cell therapy by alleviating challenges linked to apheresis-based methods, with its patient-friendly nature, cost-effectiveness, and distinctive methodology.

## INTRODUCTION

Traditionally, apheresis has been the source of cellular starting material for T-cell therapy products due to the large numbers of T-cells required to go through a more conventional manufacturing process and generate sufficient modified T-cells for a therapeutic dose to treat oncology patients. The associated burden on patients due to the length and invasiveness of the apheresis cell collection procedure and the logistical constraints of transporting the apheresis product to the manufacturing location are challenges currently associated with CAR T-cell products that limit accessibility and necessitate a new approach.

The challenges encountered in obtaining an optimal leukapheresis starting material for CAR T-cell manufacturing encompass operational hurdles like access to specialized apheresis centers that meet guidelines along with resourced and trained staff,^2^ and technical issues such as vascular access, contamination from other cell types, and effectively managing adverse events during the collection process.^3^

Furthermore, conventional CAR T-cell manufacturing involves a protracted culture of patient apheresis material over 7–10 days designed to maximize expansion, resulting in a final product with a highly differentiated T-cell phenotype. However, studies in oncology showed that T cells with younger, more stem-like phenotype are correlated with potentially improved clinical benefit compared with those with differentiated memory, effector function, or exhaustion phenotypes, which are signatures of more differentiated T-cell types. Shorter manufacturing processes can alleviate these issues and lead to improved CAR T-cell products.^4^

Starting from a whole blood draw instead of apheresis, combined with a shorter duration of CAR T-cell manufacturing, holds the potential to revolutionize the patient experience of CAR T-cell therapy by addressing key challenges associated with conventional methods. This optimization could decrease production costs,^1^ increase treatment accessibility, and increase the overall feasibility of CAR T-cell therapies.

Ingenui-T is a comprehensive CAR T-cell vein-to-vein approach initially being developed for autoimmune disease, utilizing the same fully human anti-CD19 CAR construct as KYV-101. KYV-101 is an investigational autologous anti-CD19 CAR T-cell therapy (manufactured using conventional methods) under investigation for patients with B-cell-driven autoimmune diseases, including lupus nephritis, systemic sclerosis, myasthenia gravis, multiple sclerosis, and other diseases with a strong rationale for B-cell involvement in the disease pathology. Here, we describe Ingenui-T, highlighting its ability to generate high-purity and functional CAR T-cells. Importantly, the Ingenui-T platform yields CAR T-cells with a potent functional profile and a less differentiated phenotype compared to CAR T-cells generated from a more conventional manufacturing process that uses apheresis-derived cellular starting material. By circumventing the challenges associated with apheresis, the Ingenui-T platform offers a promising solution for enhancing the efficiency and accessibility of CAR T-cell therapy, lowering costs, and ultimately advancing its application in the realm of autoimmune diseases.

## METHODS

### Patient Whole Blood and Leukapheresis Collection

Peripheral whole blood (up to 200 mL) from healthy donors (n=9, AllCells or Bloodworks Northwest, USA) was collected and transported fresh for immediate processing. Cryopreservation was intentionally avoided to maximize cell viability. A cell count was performed to quantify the incoming blood cell population. Donor-matched cryopreserved leukapheresis material (n=4, AllCells, USA) was also obtained to generate CAR T-cells according to a conventional manufacturing process.

### Ingenui-T Manufacturing Platform

Briefly, up to 200 mL of collected whole blood was added directly to the Gibco™ CTS™ DynaCellect™ Magnetic Separation System (Thermo Fisher Scientific, Waltham, MA), and Dynabeads (Thermo Fisher Scientific, Waltham, MA) were used at a defined ratio for enrichment and activation of T cells. T cells were counted, analyzed by flow cytometry for CD3^+^ T-cell purity, and seeded into vessels containing culture media enriched with cytokines. T-cells were transduced using a lentiviral vector encoding the Hu19-CD828Z anti-CD19 CAR construct at a fixed multiplicity of infection (MOI). This is the same construct used in KYV-101 - a first-in-class, fully human autologous anti-CD19 CAR T-cell therapy (Kyverna Therapeutics, Emeryville, CA). After a targeted in vitro cell culture period of less than 72 hours post-seeding, cells were harvested, formulated into final product containers, and cryopreserved **(Figure 1**). In parallel, non-transduced cells were also generated through the same manufacturing process in the absence of lentiviral transduction for use as control cells.

**Figure 1.**
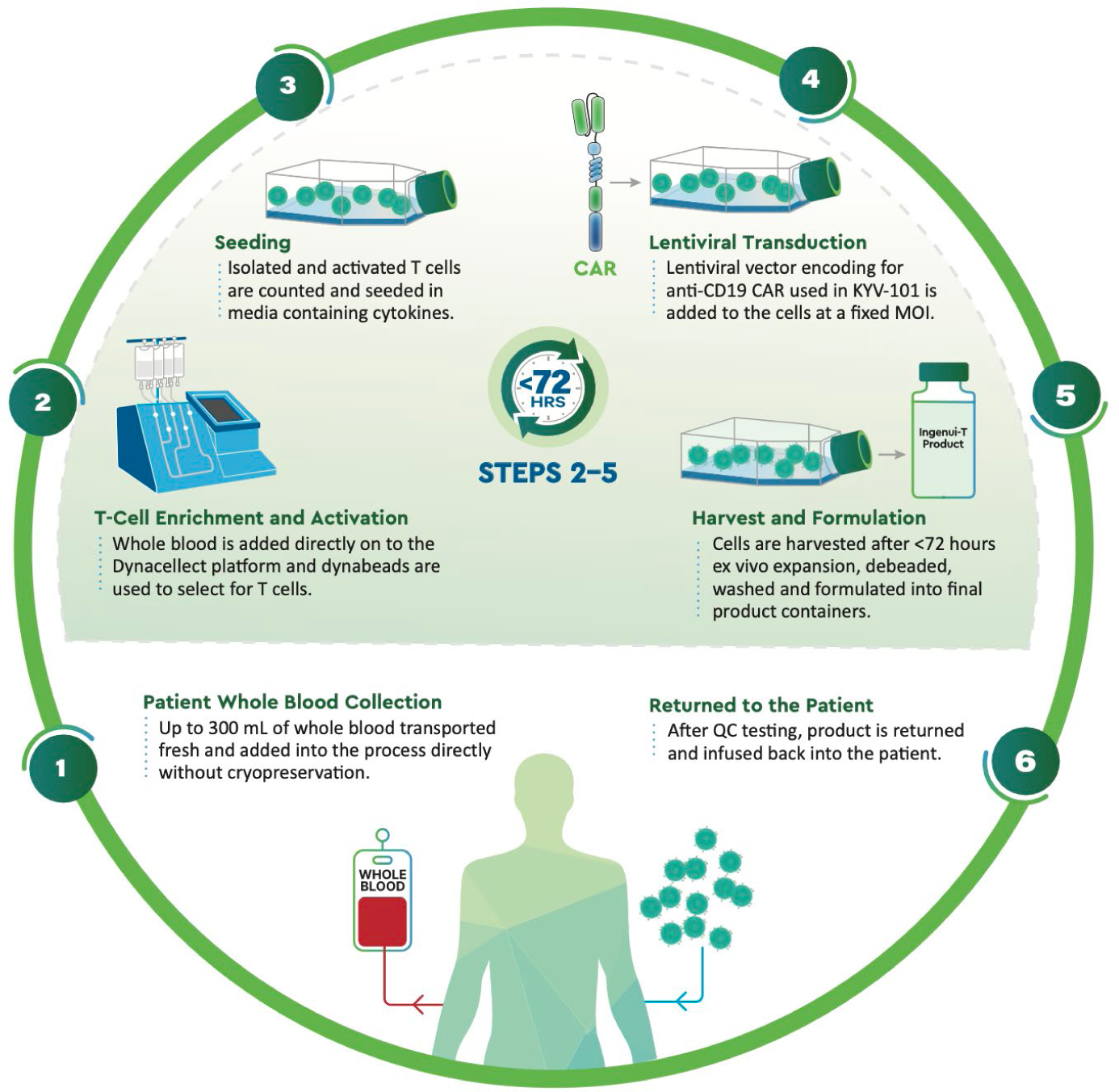
Manufacturing of CAR T-cells using Ingenui-T platform Abbreviations: CAR, chimeric antigen receptor; mL, milliliter; MOI, multiplicity of infection; QC, quality control.

### Conventional RUO CAR T-Cell Manufacturing

For conventional Research Use Only (RUO) manufacturing of anti-CD19 CAR T-cells, the cell product was manufactured to represent the KYV-101 product in the Kyverna development laboratories.

Cryopreserved leukapheresis material was thawed, washed, and subjected to antibody-driven T-cell isolation using magnetic beads (Miltenyi Biotec). The isolated T cells were counted, analyzed by flow cytometry for CD3^+^ T cell purity, and activated in the presence of media containing cytokines. As with the Ingenui-T platform, T cells were transduced at a fixed MOI using the same lentiviral vector incorporating the Hu19-CD828Z anti-CD19 CAR construct (Kyverna Therapeutics, Emeryville, CA). Cells were then cultured for >8 days before harvesting, formulation, and cryopreservation of the final product.

### In Vitro CAR T-Cell Phenotyping

Flow cytometry was used to analyze CD3^+^ T-cell purity, CD4^+^ and CD8^+^ T-cell populations, and anti-CD19 CAR expression. CD4^+^ and CD8^+^ T-cell memory phenotypes of the Ingenui-T final product were compared to the T-cell memory populations in whole blood starting material by flow cytometry analysis. Similarly, the T-cell memory phenotypes in conventional CAR T cells were compared to the T-cell memory populations in the apheresis starting material (donor-matched with whole blood Ingenui-T cells). CAR expression was analyzed at 0 hour and 72 hours post-thaw of the final drug product.

### In Vitro CAR T-Cell Functional Activity

To evaluate the functional activity of CAR T-cells in a short-term, single-challenge cytotoxicity assay, donor-matched Ingenui-T cells or conventional CAR T-cells were co-cultured with CD19^+^ target cells expressing a fluorescent protein at the indicated effector to target (E:T) ratios. Target-specific cytotoxic activity was assessed by imaging co-cultures using an Incucyte Sx5 (Sartorius) instrument and calculating the survival or outgrowth of fluorescent target cells over time.

To evaluate the long-term functionality of CAR T-cells, donor-matched Ingenui-T cells or conventional CAR T-cells were serially rechallenged every 2–3 days with CD19^+^ target cells at the indicated E:T ratios. At each time point, samples were split in half to assess the percent cytotoxicity and to re-plate with fresh target cells. Percent cytotoxicity was calculated at each timepoint by measuring the target cell survival using flow cytometry and was normalized to the survival of target cells in the absence of CAR T effector cells.

To evaluate cytolytic activity of CAR T-cells against autologous primary B cells, Ingenui-T cells or control non-transduced T cells were co-cultured with peripheral blood mononuclear cells (PBMCs) obtained from donor-matched leukapheresis material. Effector-to-target (E:T) ratios were defined according to the number of CAR^+^ Ingenui-T cells (effector) to total PBMCs (targets). After co-culture for a specified time period, target-specific cytolytic activity against CD19^+^ or CD20^+^ B cells was measured by flow cytometry. Percent cytolysis against B cells was calculated by normalizing to the survival of B cells in PBMC-only cultures.

## RESULTS

### Ingenui-T Cell Manufacturing Platform

Our objective was to demonstrate the technical feasibility of generating anti-CD19 CAR T-cells starting from fresh whole blood material in a shortened manufacturing process. As shown in **Figure 1**, fresh whole blood from healthy donors was loaded onto the DynaCellect platform using Dynabeads at a defined bead:cell ratio. The isolated T cells were sampled to confirm the isolation purity (≥95% CD3^+^) by flow cytometry and activated T cells were subsequently seeded into culture with media containing supporting cytokines. Transduction using a lentiviral vector encoding the anti-CD19 CAR construct occurred at a fixed MOI. Following a brief period in culture to allow for cell recovery and integration of the transgene (<72 hours post-seeding) CAR T-cells were collected for formulation in cryopreservation media. T-cell purity analysis of the final Ingenui-T cell product showed a T-cell percentage of 93.9±1.6%, obtained from a starting T-cell frequency of 42.3±6.8% in whole blood. These results were comparable to the T-cell enrichment obtained via a conventional CAR T-cell manufacturing process using donor-matched cells (final T-cell purity of 94.0±3.3% from a starting T-cell frequency of 46.3±7.2% in conventional apheresis; **Table 1**). Given the short culture time, Ingenui-T cells demonstrated minimal expansion during the manufacturing process, resulting in a 0.68±0.09-fold change of the total T-cell number from the time of culture seeding to final formulation (including any losses due to final harvest procedures). Nevertheless, the final yield of Ingenui-T cell product was 38.5±6.6×10^6^ T cells per 100 mL of starting whole blood. Product attributes were tested upon harvest and after 72 hours of post-thaw culture to simulate product performance in the patient. At 72 hours post-thaw of the drug product, CAR^+^ expression ranged between 45.1%–54.5% in Ingenui-T cells, which was statistically similar to the 37.4%– 56.3% CAR^+^ expression obtained from the conventional CAR T-cell manufacturing process derived from apheresis (**Table 1**). This demonstrates that using the novel Ingenui-T approach, anti-CD19 CAR T cells can be successfully manufactured directly from whole blood, in a shortened manufacturing process, at a scale sufficient for therapeutic dosing of B-cell-driven autoimmune disease patients.

**Table 1.** CAR T-cell data in conventional versus Ingenui-T manufacturing RUO processes.

| Characteristic | Conventional CAR T, from apheresis (n=4 HD) | Ingenui-T, from whole blood (n=4 HD) |
| --- | --- | --- |
| % T-cell purity in starting material | 46.3 ± 7.2 | 42.3 ± 6.8 |
| % T-cell purity in drug product | 94.0 ± 3.3 | 93.9 ± 1.6 |
| % CAR<sup>+</sup> of T-cells in drug product<sup>a</sup> | 46.5 ± 5.0 (range 37.4–56.3) | 49.3 ± 1.4 (range 45.1–54.5) |
| Fold expansion of T-cells from starting material | 28.8 ± 1.9 | 0.68 ± 0.09<sup>b</sup> |

Abbreviations: CAR, chimeric antigen receptor; HD, healthy donor (matched apheresis and whole blood); RUO, research use only.
<sup>a</sup>Analyzed at 72 hours post-thawing of drug product.
<sup>b</sup>Data compiled from n=9 donors.

### Phenotypic and Functional Comparison of Ingenui-T Cells and Conventional CAR T-Cells

To demonstrate the pharmacologic activity of anti-CD19 CAR T-cells generated in the Ingenui-T platform, we performed a set of phenotypic and functional in vitro characterization experiments, comparing Ingenui-T cells with CAR T-cells expressing the same anti-CD19 CAR construct generated with a conventional manufacturing process. Whole blood for the Ingenui-T process and apheresis for the conventional process were sourced from the same donors as part of the same collection. As expected, due to the shortened culture period in the Ingenui-T platform, Ingenui-T cells comprised a less differentiated T-cell memory phenotype than CAR T-cells generated in a conventional manufacturing process. Whole blood-derived Ingenui-T cells preserved a T-cell memory phenotype that closely resembled the phenotype observed in the starting material, with slight increases in the overall effector/memory compartment (combined T_CM_, T_EM_, and T_E_ populations). From starting material to the Ingenui-T final product, the effector/memory compartment shifted from a mean of 48.8±5.6% to 69.4±4.8% within the CD4^+^ T cell fraction, and from 42.9±3.9% to 46.6±5.5% within the CD8^+^ T cell fraction, while maintaining a substantial proportion of cells within the T_N_+T_SCM_ compartment (**Figure 2A**). In contrast, CAR T-cells obtained from the traditional manufacturing process (with a more extended culture period) had mostly converted to the effector/memory compartment, shifting from a mean of 58.4±3.5% to 94.0±2.8% and from 40.1±5.2% to 86.2±4.9% within the CD4^+^ and CD8^+^ T cell fractions, respectively. Importantly, minimizing the differentiation of Ingenui-T cells in vitro is anticipated to preserve their activation potential for in vivo expansion and activity in the patient. This in turn enables the administration of a significantly lower dose for equivalent therapeutic efficacy in clinical application, which is a critical attribute for Ingenui-T considering the inherently lower starting number of T cells that can be obtained from a whole blood draw relative to apheresis.

**Figure 2.**
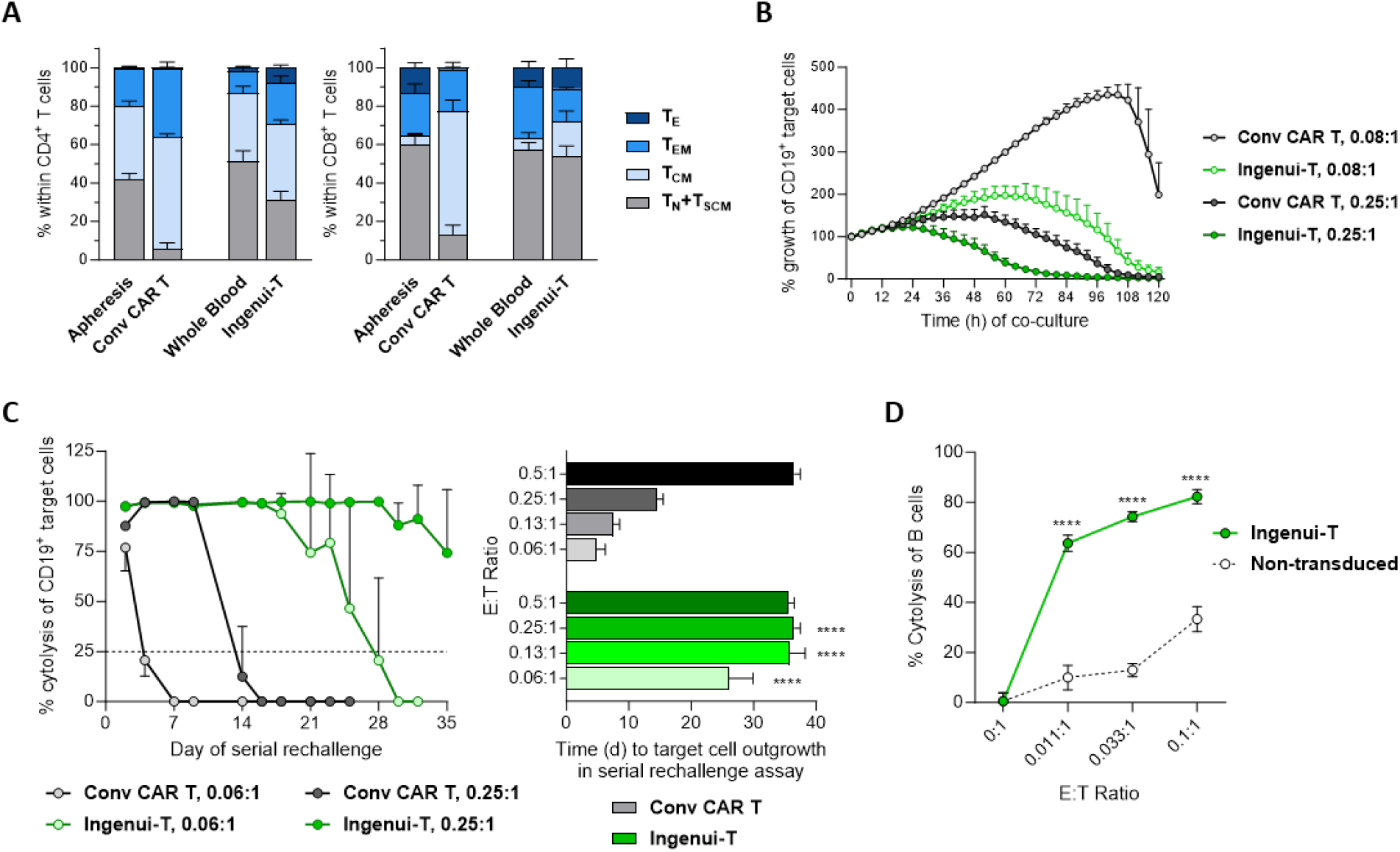
Phenotype and in vitro functionality of Ingenui-T cells Abbreviations: CAR, chimeric antigen receptor; Conv, conventional; E:T, effector to target; PBMC, peripheral blood mononuclear cell; SD, standard deviation; SEM, standard error mean; T_CM_, central memory T cells; T_E_, effector T cells; T_EM_, effector memory T cells; T_N_, naive T cells; T_SCM_, stem cell memory T cells. A) Rapid-process manufacturing of Ingenui-T better preserves the T-cell memory phenotype from whole blood starting material compared to conventional CAR T-cell manufacturing (Conv CAR T, from apheresis starting material). Data are shown as mean ± SEM, combined from n=4 healthy donors. B) Ingenui-T cells have more potent cytolytic activity than conventionally manufactured anti-CD19 CAR T-cells (Conv CAR T) in an Incucyte-based killing assay targeting CD19-expressing cells at the indicated effector to target (E:T) ratios. Data are shown as mean ± SD from 3 technical replicates per condition, from one donor representative of N=4 healthy donors. C) Ingenui-T cells have more potent and more durable cytolytic activity than conventionally manufactured anti-CD19 CAR T-cells (Conv CAR T) in a serial rechallenge killing assay. Left: Donor-matched Ingenui-T cells and Conv CAR T-cells, both derived from whole blood starting material, were co-cultured with CD19^+^ target cells at the indicated effector to target (E:T) ratios, with repeated additions of target cells every 2–3 days for up to 35 days. Survival of target cells at each timepoint was measured by flow cytometry. Data shown from one representative donor as mean ± SD from 4 technical replicates per condition. Right: Time (days) to target cell outgrowth in the serial rechallenge killing assay, indicative of the loss of effective CAR T-cell cytolytic activity, was calculated according to a decrease in target cell cytolysis below the 25% threshold (dotted line in left panel). ****p<0.0001, comparing matched E:T ratios, by 2-way ANOVA (GraphPad Prism). D) Ingenui-T cells successfully killed autologous primary B cells in a dose-dependent manner. Ingenui-T cells or non-transduced T cells, derived from whole blood, were co-cultured for a defined time period with autologous (donor-matched) PBMCs at the indicated effector to target (E:T) ratios, representing the ratio of CAR^+^ T cells (effector) to total PBMCs (target). Survival of B cells, defined by the surface expression of CD19 or CD20, was measured by flow cytometry. Data are shown as mean ± SD from 3 technical replicates per condition, from one donor representative of N=2 healthy donors. ****p<0.0001, comparing matched E:T ratios, by 2-way ANOVA (GraphPad Prism).

The functional activity of Ingenui-T cells was assessed in short-term and long-term in vitro preclinical assays in order to demonstrate target-specific cytotoxicity against CD19-expressing cells. The cytotoxic activity of Ingenui-T cells and donor-matched, apheresis-derived conventional CAR T-cells (from a conventional expansion culture, expressing the same CAR construct) were compared in a short-term cytotoxicity assay against CD19^+^ target cells. As shown in **Figure 2B**, in an Incucyte imaging-based assay, Ingenui-T cells controlled the outgrowth of target cells, more effectively than conventional CAR T-cells at the same E:T ratio. This reflects the expected increase in CAR T-cell potency and target-mediated CAR T-cell proliferation due to the less differentiated memory phenotype of Ingenui-T cells. No cytotoxic activity was observed in control non-transduced T-cells (i.e., without CAR expression; data not shown). These results confirmed the anti-CD19 target-specific activity of Ingenui-T cells and their increased functional potency relative to conventional CAR T-cells that had been generated in a conventional manufacturing process.

To further evaluate the functional activity of Ingenui-T cells, we performed a long-term serial rechallenge assay in vitro. In this assay, CAR T cells and CD19^+^ target cells were co-cultured at the indicated E:T ratios starting at day 0, followed by the serial addition of the same number of target cells every 2 or 3 days, to evaluate the potency and durability of target-specific serial cytotoxicity over an extended time. As shown in **Figure 2C**, Ingenui-T cells continued to kill target cells for a significantly longer period at a given E:T ratio and required a >4-fold lower E:T ratio than donor-matched CAR T-cells generated from a conventional expansion culture to maintain the same duration of killing. This finding again matched our expectation that the younger differentiation phenotype of Ingenui-T cells results in higher functional potency, greater proliferation (data not shown), and more prolonged cytolytic activity compared to conventional CAR T-cells, which are more prone to exhaustion and loss of functionality over time.

Finally, we assessed the in vitro cytolytic activity of Ingenui-T cells against autologous primary B cells, which are the cells targeted for depletion in the treatment of patients with B-cell-driven autoimmune diseases. When Ingenui-T cells were co-cultured with autologous total peripheral blood mononuclear cells (PBMCs), B cells were eliminated in a specific and dose-dependent manner (**Figure 2D**). As expected, Ingenui-T cells demonstrated greater potency of B cell killing than conventional CAR T-cells when tested at dose-limiting E:T ratios (e.g., at 0.011:1; **Figure 2D** and data not shown). These preclinical assays provide proof-of-concept data that highly functional anti-CD19 CAR T-cells could be generated using our Ingenui-T platform starting from fresh autologous whole blood material. These results have paved the way for the clinical development of Ingenui-T cells as a therapy for autoimmune disease, which will improve the patient experience, increase treatment accessibility, and reduce costs, by eliminating the need for patients to undergo apheresis.

## DISCUSSION

Ingenui-T is a comprehensive vein-to-vein solution, focused on enhancing patient experience and reducing the cost of manufacturing CAR T-cell therapies. The next-generation manufacturing process starts from autologous whole blood and uses a rapid (<3 day) manufacturing process, resulting in a potent CAR T-cell product with demonstrated target-specific killing activity. This manufacturing process marks a significant departure from traditional methods that necessitate apheresis, a laborious and resource-intensive process, and extended cell culture.^3^

The use of whole blood eliminates the need for apheresis, alleviating the burden on the patient as it reduces the need for a lengthy and invasive collection process, ultimately enhancing the overall patient experience. Further, there is a fixed quantity of beds available for apheresis which limits the number of patients who can be treated, and which is becoming an even bigger concern as CAR T-cell therapies extend to non-oncology indications such as autoimmunity where patient numbers are significantly higher (millions of patients compared to 10s of thousands). Ingenui-T reduces the differentiation of the cells in vitro by minimizing the culture time, resulting in both a shorter process and a more potent product that can potentially provide equivalent therapeutic benefit with a lower dose, while enabling the feasibility of using a limited volume of whole blood rather than apheresis as starting material. Streamlining the process through a combination of collecting up to 300 mL of whole blood, and minimizing the culture time, not only reduces and optimizes resource utilization, but also reduces the time spent in specialized facilities and reduces involvement of highly skilled personnel, enhancing the cost-efficiency of CAR T-cell therapy. This reduction in the overall cost of goods to manufacture and decreased burden on patients holds promise for broader accessibility and affordability.^1^ This optimization also aligns with the goal of scalability of CAR T-cell therapies, addressing a critical need in the field.

The results of the innovative Ingenui-T approach demonstrate the ability to enrich T cells from whole blood and successfully generate potent anti-CD19 CAR T-cells, comparable or superior in activity to anti-CD19 CAR T-cells derived from apheresis material in a conventional manufacturing process. Phenotypic characterization indicates a less differentiated phenotype in CAR T-cells made using Ingenui-T compared to conventionally manufactured CAR T-cells. This characteristic may have clinical implications, as less differentiated T cells are associated with enhanced in vivo expansion and efficacy.^3^

Ingenui-T introduces a novel paradigm with a vein-to-vein approach for treating patients with B-cell-driven autoimmune diseases with CAR T-cell therapy, by addressing key challenges associated with traditional apheresis-based manufacturing methods. The reduced burden on patients and potential for cost-efficiency highlight Ingenui-T’s potential to significantly impact the feasibility and accessibility of CAR T-cell therapy for autoimmune diseases.

## ACKNOWLEDGMENTS

We acknowledge Sarah Franco for support with manuscript authoring; Ramona Pufan and Jennifer Leslie with manuscript review.

